# *Wolbachia*: The architect of microbial assemblies in response to environmental changes

**DOI:** 10.1101/2025.07.03.663013

**Authors:** Pina Brinker, Joana Falcao Salles, Leo W. Beukeboom, Michael C. Fontaine

## Abstract

Translocating organisms from their habitats to laboratories, a method common in microbial ecology and evolution experiments, alters their microbial communities. However, the impact of translocation on microbiomes is often overlooked, limiting our ability to assess how laboratory findings relate to field dynamics. It’s also crucial to examine organism responses with symbionts, as they can stabilise communities or increase vulnerability to changes. This study investigates the effects of laboratory introduction on host-microbiome interactions in the parasitic wasp *Asobara japonica* and its endosymbiont *Wolbachia*. Results indicate that transferring *Wolbachia*-infected (asexual) and uninfected (sexual) wasps to a laboratory decreased *A. japonica’s* bacterial diversity. Stochastic processes drive community changes, with distinct alterations in bacterial composition between reproductive modes. Over four generations, asexual wasps’ bacterial communities became more uniform, while sexual wasps showed greater diversity. Notably, bacterial changes appeared across generations rather than immediately. Additionally, *Wolbachia* abundance changed following laboratory introduction, affecting community structure and assembly. In conclusion, our research demonstrates how laboratory conditions affect host-associated microbial communities depending on symbiont presence. This, in turn, can affect their functions, host interactions, and overall community dynamics. Thus, our findings stress the importance of linking lab results to real-world implications, posing a broader challenge for researchers.

## Introduction

The host associated microbial community, the microbiome, is an essential component of all organisms and is shaped by intricate interactions with the host, among microbial members, and environmental conditions (Leibold *et al*., 2004; Adair and Douglas, 2017; Brinker *et al*., 2019, 2023; Uren Webster *et al*., 2020). This community and its interactions are sensitive to disturbances and, if disrupted, can, for example, in insects, lead to starvation (Hosokawa *et al*., 2010), susceptibility to parasites (Dheilly *et al*., 2017), or undermine disease resistance (Dacey and Chain, 2020). Moreover, disruptions not only have the potential to alter the impact of symbionts on both the host and its associated microbial community (Bénard *et al*., 2020) but can extend to the entire ecosystem within which the host resides (Pita *et al*., 2018; Schapheer *et al*., 2021).

Disruptions of host-associated microbial communities and their interactions are often caused by environmental changes (de Vries *et al*., 2004; Russell and Moran, 2006; Ochman *et al*., 2010; Colman *et al*., 2012; Ferguson *et al*., 2018; Duan *et al*., 2020). One peculiar and substantial environmental change is the translocation of organisms from nature into the laboratory or *vice versa* (Gall *et al*., 2017; Waltmann *et al*., 2019). The translocation of an organism can lead to changes in the diversity and abundance of microbial communities associated with the organism and thus influence essential traits of the relocated host. After the translocation from the wild into the laboratory, various environmental factors experienced by a host are likely to change, with missing fluctuations in temperature, humidity, and other factors, as well as changes in diet likely having percolating effects on the host-associated microbial community (Ochman *et al*., 2010; Colman *et al*., 2012; Luo *et al*., 2021). Moreover, the translocation from the wild into the laboratory has the potential to influence the horizontal acquisition of free-living microbes present in water, soil, or air and microbes from other species, which can negatively affect hosts if they rely on environmental acquisition or horizontal transmission of microbes (Pons *et al*., 2019; Acevedo *et al*., 2021). Finally, from a host perspective, individuals may experience a decrease in population size and a reduction in genetic diversity, potentially leading to changes in the microbial community (Smith *et al*., 2015; Brinker *et al*., 2023). Translocations of organisms and concomitant environmental changes are not only relevant for studies focusing on understanding symbiotic interactions (Brinker *et al*., 2019) but additionally carry economic implications, given the increasing commercial breeding of organisms for purposes such as feed and food production (Francuski and Beukeboom, 2020) or their deployment as biological agents for pest control (Parra and Coelho, 2022) and disease management (Moreira *et al*., 2009; King *et al*., 2018). Finally, with anthropogenic activities leading to drastic environmental changes and microbiome research relying on experimental work in laboratories and the translocation of organisms from nature into the laboratory and *vice versa*, which inherently involve environmental alterations, an understanding of host-associated microbial community reactions to environmental changes during or after translocation will become more and more important.

We know that individuals reared in the laboratory harbour less diverse microbial communities compared to those living in the wild. However, the changes in microbial communities after laboratory introduction need to be evaluated to determine whether they occur randomly (stochastically) or follow a pattern (deterministically), which could inform future modelling approaches that predict such changes. Additionally, it’s important to investigate if changes occur differently in systems in which a powerful symbiont is present, as symbionts could act as a stabiliser of the microbial community (Herren and McMahon, 2018) or make them more susceptible to changes. To investigate this, we will use a simple model system represented by the haplodiploid parasitic wasp *Asobara japonica*. Like other insects, *A. japonica* has a relatively low microbial diversity (Engel and Moran, 2013; Brinker *et al*., 2023) and naturally occurs with and without the endosymbiotic bacterium *Wolbachia*, whose presence modulates host reproduction. *Wolbachia* infected wasps reproduce asexually, producing only females from unfertilised eggs (thelytokous parthenogenesis; (Kremer *et al*., 2009)), whereas uninfected wasps reproduce sexually (arrhenotoky). A comparison between infected and uninfected wasp lines thus allows us to infer whether the presence of symbionts influences patterns of microbial community change associated with translocations through microbe-microbe interactions (Brinker *et al*., 2019). In addition, the use of three sexual (uninfected) and three asexual (infected) distinct genetic lines of *A. japonica* (Brinker *et al*., 2023) allows us to infer whether patterns depend on host genetic background, population structure, and microbiome composition. We first evaluate changes in the wasps’ associated bacterial communities across four generations after their transfer from nature to the laboratory. We anticipate changes in both infected and uninfected wasps, with the most significant shifts in the initial generations due to substantial environmental changes after translocation. We also explore the influence of *Wolbachia* on these changes, hypothesising that infected (asexual) wasps will exhibit fewer microbial community changes, thanks to potential community-stabilising effects (Herren and McMahon, 2018) and a lack of genetic diversity loss from asexual reproduction. In contrast, sexual wasps not only lack a potential stabilising key bacterial taxon but also experience a reduced pool of potential mating partners, causing a bottleneck effect leading to a loss of genetic variability. Therefore, we expect the microbial community changes of sexual wasps to be more substantial than the ones of asexual wasps. Lastly, we investigate whether deterministic or stochastic processes govern bacterial community changes upon laboratory introduction, aiming to discern predictability patterns in these shifts.

## Material and methods

### Wasp collection and rearing

*Asobara japonica,* a larval parasitoid of various *Drosophila* species native to southeastern Asia, occurs naturally infected with *Wolbachia,* causing asexual (thelytokous parthenogenetic) reproduction on Japan’s main island and uninfected, reproducing sexually on the southern islands of Japan (Mitsui *et al*., 2007). For this study, wasps from six locations (three infected with *Wolbachia*, three uninfected; see Fig. 1) were collected in June 2017. Each of these locations represents a genetically distinct population (Sexual Pop 1 – Pop 3, Asexual Pop 4), except the two most northern ones, Kyoto and Sendai, which belong to the same population (Pop 5) (Brinker *et al*., 2023). Asexual populations occur in different environmental conditions, with subtropical conditions in Kagoshima (Pop 4) and more temperate conditions in Kyoto and Sendai (both Pop 5). In contrast, all sexual locations have a subtropical climate.

**Figure 1:**
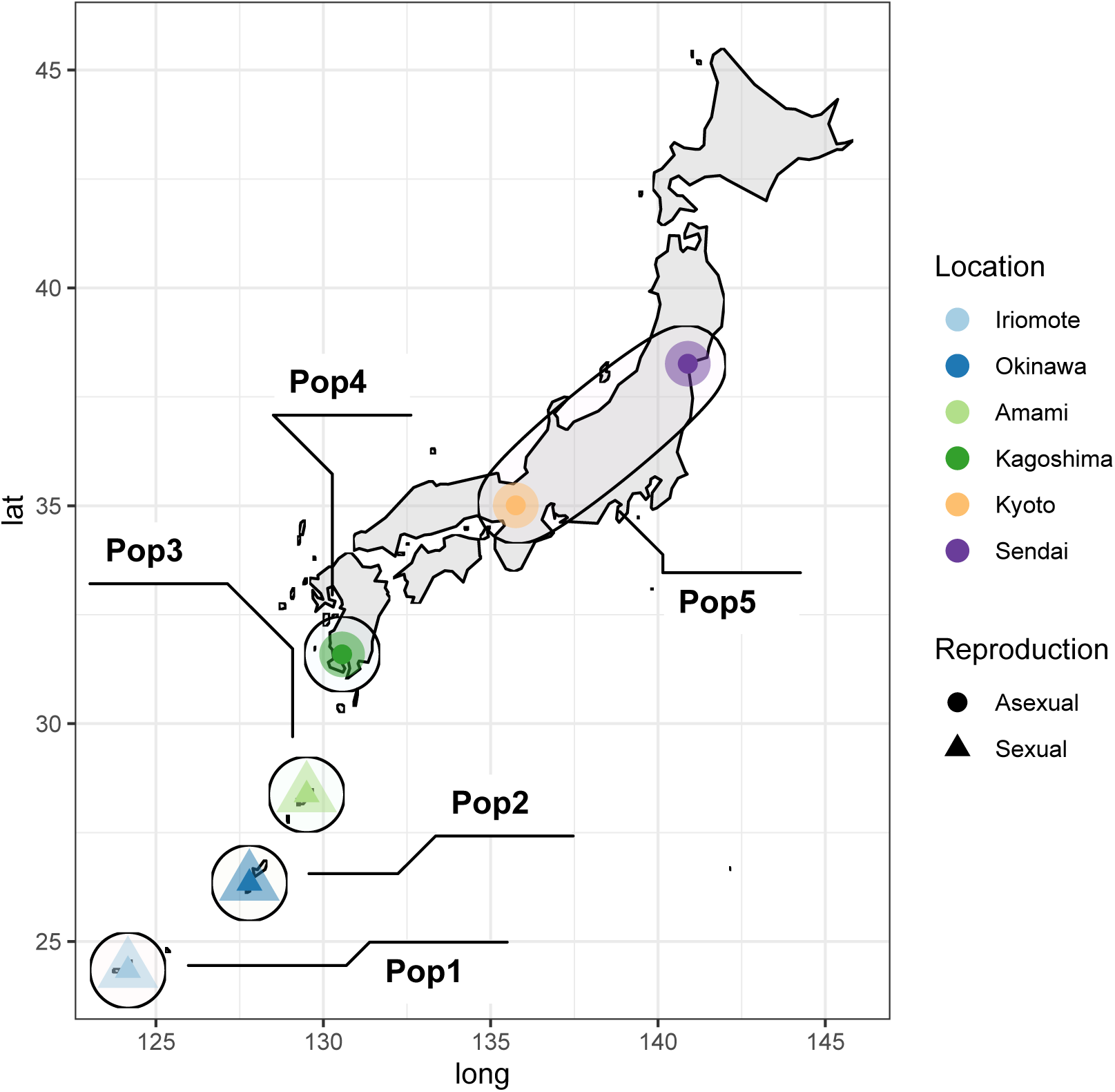
Schematic map of Japan showing collection sites of *Asobara japonica* used to establish laboratory lines in this study. Locations with sexually reproducing wasps in the south of Japan are indicated by a triangle and locations with asexually reproducing wasps in the north of Japan by a circle. Black ellipses show the population identity of wasps collected from the location inferred from population genetic analyses (see Brinker *et al*. 2023).

Wasps were collected as larvae and brought to the laboratory in the Netherlands, where adults hatched. This way, we ensured a natural starting point for the subsequent laboratory rearing. From these G0 mother wasps, we created seven replicate iso-female lines from each location. Lines were maintained for another four generations in stable laboratory conditions (25°C, LD16:08 light cycle). Each new generation was started by placing three to four females (offspring of the previous generation, matured for three days during which they were fed honey) into agar bottles coated with a thin layer of yeast (AB Mauri S.p.A., Italy) and containing second instar *D. melanogaster* (ww-strain) larvae. For the sexually reproducing wasps, two to three males were added to each bottle to secure mating of females. Ten females per replicate were frozen at −80°C for microbial DNA extraction in each generation, resulting in seven coherent replicates for each location and generation.

### Bacterial DNA extraction

For bacterial DNA extraction, ten pooled wasps were first washed to remove any environmental contamination (1min in 70% ethanol and 3x in sterile water). DNA was then extracted and purified using the DNeasy Power Soil^®^ DNA Isolation Kit, following the manufacturer’s protocol (Power Soil^®^, MoBio Laboratories Inc., California, United States), except that wasps were snap-frozen in liquid nitrogen and crushed with a sterile pestle in a 1.5ml tube before being added to the homogenisation tubes and homogenised for 15min using a grinder (Kaiser, Germany). After extraction, DNA was eluted in 100µl C6 solution provided by the manufacturer and stored at −20°C for further processing and Illumina MiSeq sequencing (2×300bp MiSeq, ∼40,000 reads/sample) of the V4 region of the bacterial 16S rRNA gene to the Minnesota genomic centre (Gohl *et al*., 2016).

### Bacterial community analyses

Demultiplexed 300bp paired-end reads were processed following the *dada2* pipeline v.1.18.0 (Callahan *et al*., 2016) in *R* v. 4.0.2 (R Core Team, 2020). First, read quality was checked by visualising the quality score. Positions with a lower mean quality score than 30 were removed. The first ten nucleotides of forward and reverse reads were trimmed, and the forward reads truncated to 250, and the reverse reads to 230 bases. Next, reads were dereplicated, and expected errors were removed. This was followed by merging of the reads and removal of chimaera errors. Finally, taxonomy was assigned using the pre-trained Silva 138 taxonomy classifier (McLaren and Callahan, 2021), creating an amplicon sequence variants (ASV) table and an unrooted neighbour-joining tree using the *phangorn* v.2.7.1 R package (Schliep, 2011; Schliep *et al*., 2017). These output files were then used to create a phyloseq object (R package *phyloseq* v.1.34; (McMurdie and Holmes, 2013)). Sequences identified as chloroplast, mitochondria, archaea, or uncharacterised reads at the phylum level, and three outliers based on DNA quality (Sexual: A24 G4; Asexual: Ky2 G2; Ky19 G3) were removed from this object. The dataset was rarefied to 1196 reads and will be referred hereafter as the “full dataset”. From the full dataset, a second phyloseq object was created in which reads belonging to the genus *Wolbachia* were removed. Taxa abundance of the phyloseq object without *Wolbachia* reads was normalised following the *edgeR* method (*microbiomeSeq* v.0.1 R package; (Ssekagiri, 2020)) prior to ordination analyses and will be referred to as “reduced dataset” hereafter. Moreover, to investigate the effect of laboratory introduction on the total endosymbiont itself, *Wolbachia* reads were extracted from the normalised (*edgeR* method, *microbiomeSeq* v.0.1 R package; (Ssekagiri, 2020)) unrarefied dataset to create a *Wolbachia* read count table.

Statistical analyses were performed in *R* v.4.0.2 (R Core Team, 2020). All analyses comparing patterns in asexual *vs* sexual lines were run with the full and reduced datasets to infer the effect of the high abundance of *Wolbachia* reads in the asexual samples on the analyses. Alpha diversity estimates (i.e., observed number of amplicon sequence variants (ASV) and Shannon diversity) were calculated with the *phyloseq* function “*estimate_richness*”. Diversity estimates were analysed in separate linear mixed-effects models (LMM, R package “*lme4*”; (Bates *et al*., 2015)) with generation (G1 to G4), reproductive mode (asexual, sexual), and their interactions as fixed predictors and location as a random effect. Similarly, the *Wolbachia* read counts were analysed in separate linear mixed-effects models (LMM, package “*lme4*”; (Bates *et al*., 2015)) with generation (G1 to G4) as fixed predictor and location as a random effect. To assess the significance of the predictor generation, this model was compared to a null (intercept only) using a likelihood ratio test (LRT). To assess the significance of predictors, models were compared to null (intercept only) or reduced models using likelihood ratio tests (LRT). Model assumptions were checked with model diagnostic tests and plots implemented in the package *DHARMa* v.0.4.4 (Hartig, 2021). As reproductive mode depended on generation for observed ASV number (significant interaction), we created two subsets separating asexual and sexual data for further testing. Pairwise comparisons between factor levels of a significant predictor in these models were performed using Tukey post-hoc tests adjusting the family-wise error rate according to the method of Westfall (package *multcomp*; (Hothorn *et al*., 2008)). Additionally, the effect of generation on observed ASV number and Shannon diversity, as well as the effect of generation on *Wolbachia* read counts, was analysed separately for each location using an ANOVA followed by pairwise comparisons for the reproductive modes. This was done considering each location as a different host genetic background, as each location is a genetically distinct population (except Kyoto and Sendai, which belong to the same population).

Variation in community composition (beta diversity) among samples was visualised via a principal coordinates analysis (PcoA) based on the Bray–Curtis distance dissimilarity matrices created with the phyloseq function “ordinate”. The number of significant PCo dimensions to interpret was determined based on a visual examination of the scree-plot (Fig. S1, S2). The effect of reproductive mode (asexual, sexual), lab rearing over the four generations, and location identity, *a.k.a.* genetic background, were tested by performing permutational multivariate analysis of variance using the function “adonis” and 999 permutations (PERMANOVA) using the v*egan* R package; (Anderson and Willis, 2003; Oksanen *et al*., 2016).

Lastly, we assessed whether bacterial community assembly is governed by deterministic or stochastic processes, or both and whether these processes are influenced by reproductive mode. For this, we followed the methods outlined in Stegen *et al*. (2012, 2013). The pairwise phylogenetic turnover between communities was calculated by the mean nearest taxon distance metric (βMNTD) using the “*comdistnt*” function (abundance.weighted = TRUE) in “*picante*” (Kembel *et al*., 2010). The difference between the observed βMNTD and the mean of the null distribution was measured in units of standard deviations (of the null distribution), using the convention to β-Nearest Taxon Index (βNTI) (Stegen *et al*., 2013). βNTI values (derived from pairwise comparisons) were compared between the different generations for both reproductive modes. In brief, βNTI *<* −2 or *>* +2 indicates that βMNTD_obs_ deviates from the mean βMNTD_null_ by more than two standard deviations. Thus, βNTI *<* −2 or *>* +2 deviate significantly from the model-based expected phylogenetic turnover, and thus turnover is driven by deterministic processes. Here, βNTI *<* −2 indicates low turnover (i.e., homogeneous selection), and βNTI *>* 2 indicates high turnover (i.e., variable selection). Values between −2 and +2 indicate the lack of deviation, implying the dominance of stochastic processes, where community assembly is less influenced by selection and more by chance (Stegen *et al*., 2012; Dini-Andreote *et al*., 2015). Additionally, the Bray–Curtis-based Raup–Crick metric (Rcbray) was calculated (Stegen *et al*., 2012, 2013) to partition further the relative influences of stochastic processes. Rcbray > 0.95 indicates dispersal limitation, meaning species movement is restricted, leading to higher community dissimilarity (Zhou and Ning, 2017). Rcbray < −0.95 represents homogenising dispersal, meaning high species movement, resulting in decreased community dissimilarity (Zhou and Ning, 2017). Rcbray values between −0.95 and 0.95 suggest undominated processes when neither selection nor dispersal dominates community assembly (Dini-Andreote *et al*., 2015).

## Results

### Bacterial diversity changes over generations

Bacterial community diversity in *Asobara japonica* was assessed across three sites for sexual wasps and three sites for *Wolbachia*-infected asexual wasps over four generations of laboratory culturing. A total of 1,203 microbial taxa were identified among 3,779,732 sequence reads. After rarefaction to 1,196 reads per sample, 827 taxa were retained, comprising a total of 188,968 reads. Five samples were excluded because they fell below the read threshold, with one in Kyoto (generation two), two in Iriomote (generation two), one in Iriomote (generation four) and one in Okinawa (generation four). Upon removing “*Wolbachia*” reads, the dataset contained 819 taxa and 114,302 reads.

Sexual wasps in which *Wolbachia* is absent exhibited a higher species diversity than the asexual wasps. However, in both sexual and asexual wasps, alpha diversity decreased over the four generations of laboratory culturing (Fig. 2). This decrease was dependent on the reproductive mode for the number of bacterial species (LMM; interaction generation x reproductive mode; LRT, χ^2^=10.94, df=3, *p*=0.012), but not for Shannon diversity (LMM; interaction generation x reproductive mode; LRT, χ^2^=5.92, df=3, *p*=0.116). To delve deeper into the effect of alpha diversity changes (observed ASV number and Shannon diversity) across generations, we separated the data into the two reproductive modes: sexual and asexual lines. For both reproductive modes, observed ASV number and Shannon diversity were significantly affected by generation (Fig. 2; sexual lines: observed ASV number: LRT, χ^2^=18.42, df=3, p<0.001; Shannon: LR-Test, χ^2^=19.28, df=3, p<0.001; asexual lines: observed ASV number: LR-Test, χ^2^=18.31, df=3, p<0.001; Shannon: LR-Test, χ^2^=19.66, df=3, p<0.001). However, there were differences in the rate of change over generations between sexual and asexual lines. Sexual lines showed a reduction in alpha diversity already after the first generation (Fig. 2, Table S1). In contrast, asexual lines showed a reduction in diversity only after the second generation (Fig. 2, Table S1). Further examination of this generation effect for each wasp line revealed that these patterns were driven by one line for both reproductive modes: Okinawa for sexual and Kagoshima for asexual reproducing wasps, respectively (see Fig. S3, Table S1). The Okinawa line (Pop 2) showed a significant decrease in alpha diversity (ANOVA: observed ASV number F_3,21_=10.26, p<0.001; Shannon F_3,21_=4.582, p=0.013) after the first generation and the Kagoshima line (Pop 4), at generation 3 (ANOVA: observed ASV number F_3,24_=7.72, p<0.001; Shannon F_3,24_=25.86, p<0.001; Table S1).

**Figure 2:**
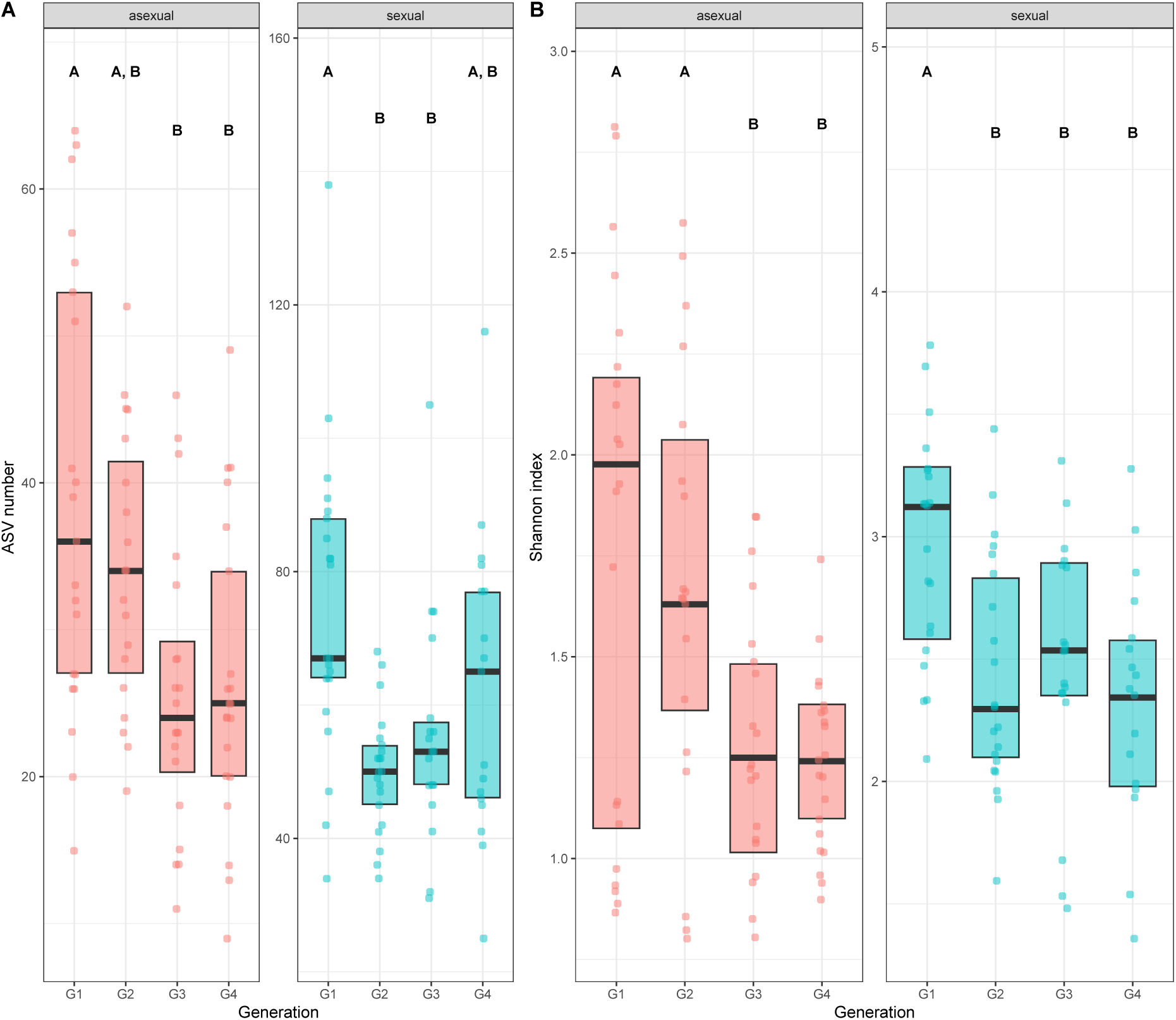
Alpha diversity expressed as **A)** ASV number and **B)** Shannon Index of asexual (*Wolbachia* infected) and sexual (*Wolbachia* uninfected) *A. japonica* wasps reared in seven replicated lines over four generations (G1 to G4) in the laboratory. Boxplots show the median and interquartile range. Letters above boxes indicate significant differences between generations.

Removal of *Wolbachia* reads from the dataset did not change these patterns (Fig. S4, Table S1), with the exception that the pattern for Shannon diversity changed in asexual wasps. It became more similar to the pattern observed in sexual wasps, i.e., a reduction occurred already after the first generation. The reduced dataset revealed a significant interaction between generation and reproductive mode for observed ASV number (LRT, χ^2^=10.74, df=3, p=0.013) but not for Shannon diversity (LRT, χ^2^=0.42, df=3, p=0.74). To maintain comparability between the full and reduced datasets, we again examined the effect of alpha diversity changes (observed ASV number and Shannon diversity) over generations separately for each reproductive mode. For sexual wasps, observed ASV number and Shannon diversity were significantly influenced by generation (Fig. S4; Observed ASV number: LR-Test, χ^2^=19.049, df=3, p<0.001; Shannon: LR-Test, χ^2^=20.05, df=3, p<0.001), with generation one differing significantly from the other three generations (Fig. S4, Table S1). Also, in asexual wasps, observed ASV number and Shannon diversity were still significantly influenced by generation (Fig. S4; observed ASV number: LR-Test, χ^2^=18.14, df=3, p<0.001; Shannon diversity: LR-Test, χ^2^=13.2, df=3, p<0.001), with a significant decrease occurring in generation three (Fig. S4, Table S1). Investigating this generation effect for each location individually and testing the influence of genetic background, we found that these patterns were driven by two lines, Kagoshima (Pop 4) and Kyoto (Pop 5). The asexual lines showed a significant reduction in observed ASV number (ANOVA: Kagoshima: observed ASV number F_3,24_=7.67, p<0.001; Kyoto: observed ASV number F_3,24_=3.35, p=0.003) after the second generation (Fig. S5, Table S1). A significant reduction of Shannon diversity was only found for wasps from Kyoto (ANOVA: Shannon F_3,24_=3.79, p=0.02), which occurred again at generation three (Fig. S5, Table S1). Removal of *Wolbachia* reads also revealed that asexual wasps became more similar in Shannon diversity, indicating that *Wolbachia* is one of the most significant changing taxa in the asexual lines after laboratory introduction.

### Changes in bacterial community composition differ between reproductive modes

Similar to species diversity, we found distinct differences in the community composition between *Wolbachia* infected (asexual) and uninfected wasps (sexual) (adonis: pseudo-F_1,156_=96.67, R^2^=0.38, p=0.001), with reproductive mode, explaining 40.5% of the variation and the PCoA showing a clear separation of the sexual and asexual reproducing lines (Fig. 3). Moreover, generation explained 11.3% of the variation in the data (adonis: pseudo-F_3,154_=2.69, R^2^=2.69, p=0.003). The community diversity of asexual wasps became more similar over generations, whereas that of sexual wasps became more distinct, with generation one having the lowest dispersion in sexual lines and asexual lines forming a tight cluster in generations three and four (Fig. 3). Considering the genetic background over generations, sexual wasps from Amami (Pop 2) and Okinawa (Pop 3) clustered together. In contrast, the bacterial community composition of sexual wasps from Iriomote (Pop 1) became more variable over generations (Fig. S5). For the asexual wasps, lines from Kagoshima (Pop 4) in generations one and two and Kyoto (Pop 5) in generation 1 formed distinct clusters from the other asexual wasps (Fig. S5; adonis: pseudo-F_5,152_=23.61, R^2^=0.44, p=0.001). The reduced dataset, i.e., where *Wolbachia* was removed, revealed a similar pattern. Although the clustering for the two reproductive modes became less distinct, it still explained 19% of the variation (adonis: pseudo-F_1,156_=12.49, R^2^=0.074, p=0.001). Again, generation influenced clustering (adonis: pseudo-F_3,154_=3.34, R^2^=0.06, p=0.001), explaining 14.1% of the variation. Asexual lines grouped closer together over time, whereas sexual lines became more distinct from each other over generations (Fig. S6). Looking at the patterns for single lines, we found that Kagoshima (Pop 4) and Kyoto (Pop 5) had a broader dispersion in the first two generations and became more similar to the community of Sendai (Pop 5), in which the bacterial community composition showed little variation over time (Fig. S7). The bacterial diversity of sexual wasps depicted a different pattern and became more diverse with each generation, especially the location Iriomote (Fig. S5, Fig. S7; adonis: pseudo-F_5,152_=5.64, R^2^=0.16, p=0.001).

**Figure 3:**
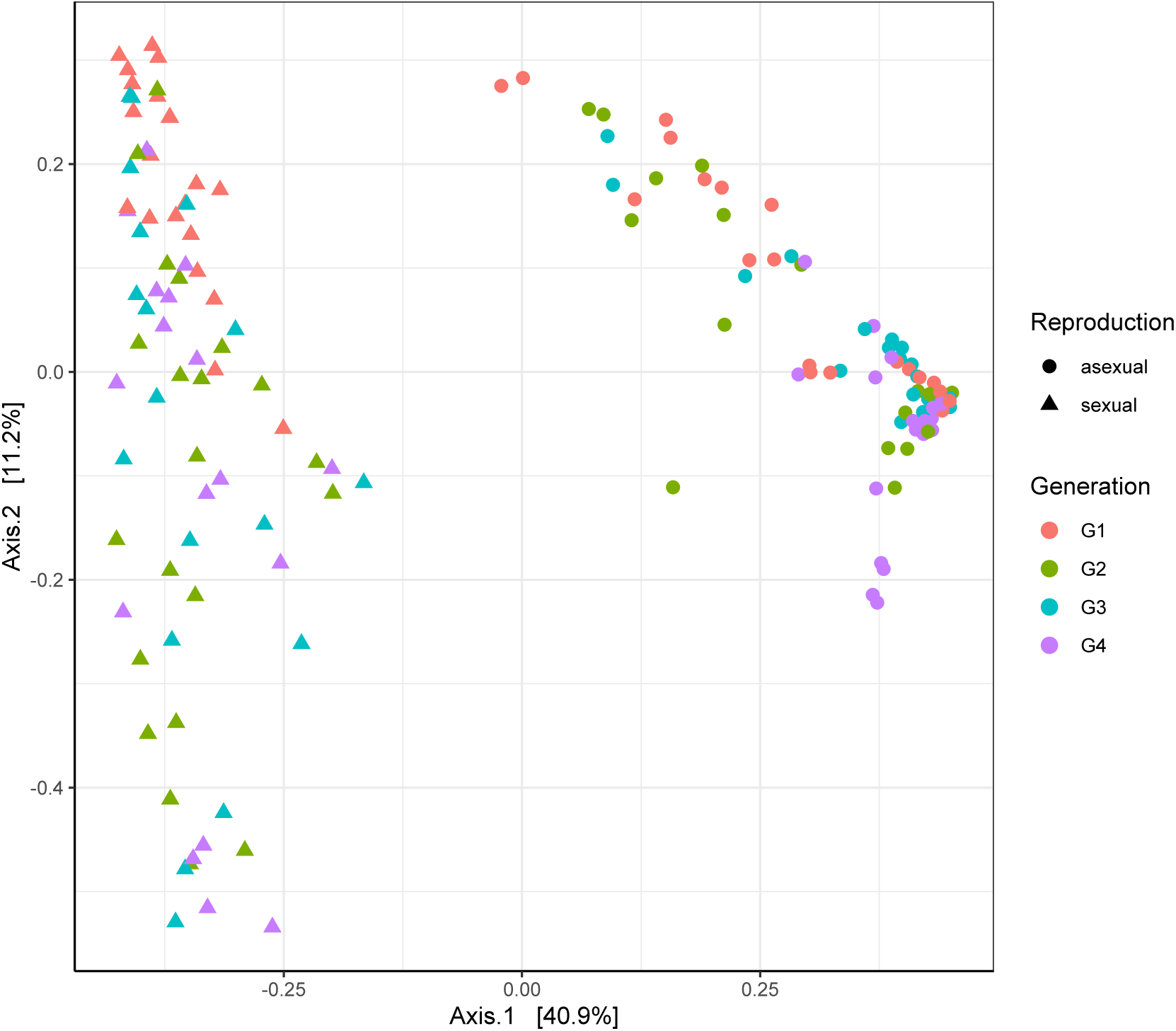
Differences in bacterial community composition between *Wolbachia*-uninfected, sexual lines (dots) and *Wolbachia*-infected, asexual lines (triangles) of *A. japonica* over four generations (colours) visualised via a principal coordinates analysis (PCoA) based on Bray– Curtis distance dissimilarity matrices.

### Number of *Wolbachia* reads changes over generations

We observed a significant fluctuation of *Wolbachia* over generations (Fig. 4A; LR-Test, χ^2^=7.395*10^−5^, df=3, p<0.001) with a decrease of endosymbiont abundance at generation 2 (mean *Wolbachia* reads ± S.D in Table S2), followed by an increase in generation three to levels similar to generation one, which then stabilised in generation four (mean *Wolbachia* reads ± S.D in Table S2; post-hoc pairwise comparisons: G2 vs G3 and G4: all p<0.001; G1 vs G3 and G4, G4 vs G3: all p>0.367). These fluctuations of *Wolbachia* do not correlate with the changes in Shannon diversity and suggest causality between the reduction of *Wolbachia* in generation two and the decrease of Shannon diversity in generation three (Fig. 4B). Moreover, the fluctuations of *Wolbachia* correlate with the homogenisation of the bacterial community composition (beta diversity) in generation three and four in asexual wasps (Fig. 3). Investigating the three asexual locations separately, we find that wasps from lines with an initially high number of *Wolbachia* (Pop 5, Kyoto and Sendai) experienced a decrease in the second generation (Fig. S8A; mean *Wolbachia* reads ± S.D in Table S3; Kyoto ANOVA: F_3,22_=4.996, p=0.009, post-hoc pairwise comparisons: G1 vs G2, G2 vs G3 and G4: all p<0.026; all other comparisons: ns; Sendai: ANOVA: F_3,24_=2.2, p=0.114). In contrast, Kagoshima (Pop 4), which had a comparably lower count of *Wolbachia* in the first generation, showed an increase of *Wolbachia* after the second generation (Fig. S8B, mean *Wolbachia* reads ± S.D in Table S3; ANOVA: F_3,24_=4.318, p=0.014, post-hoc pairwise comparisons: all p>0.065).

**Figure 4:**
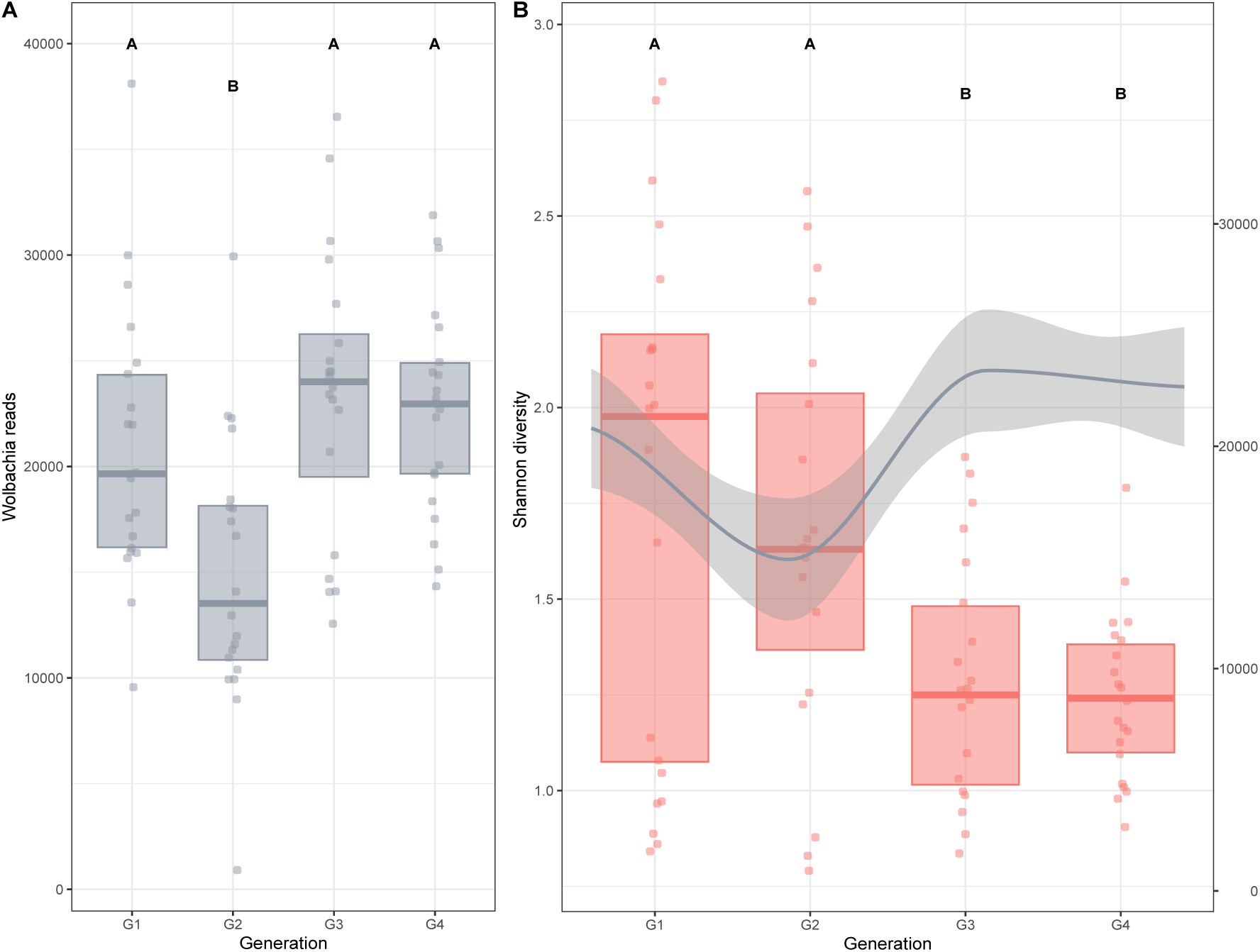
A) Abundance of *Wolbachia* cells measured as the number of *Wolbachia* sequenced reads in seven replicate lines of three asexual, *Wolbachia*-infected *A. japonica* populations (Kagoshima, Kyoto, Sendai) over four generations (G1 to G4) in the laboratory. Reads extracted out of the unrarefied but normalised full dataset are plotted. Boxplots show the median and interquartile range. **B)** Plot of the Shannon diversity (red boxplots, left y-axis) and a LOESS-smoother (locally weighted running line smoother) together with confidence intervals on *Wolbachia* reads (grey line and grey area show 95% confidence intervals, right y-axis). Letters above boxes denote statistically significant differences between generations for *Wolbachia* sequence reads (**A**) and Shannon diversity (**B**).

### Stochastic processes govern bacterial community assembly

Applying the framework of Stegen et al. (2012, 2013), we found that bacterial community assembly over the four generations was mainly governed by stochastic processes (Fig. 5), such as dispersal limitation −constraint movement of species leading to higher community dissimilarities-, homogenising dispersal −high levels species movement lead to more similar communities- (Zhou and Ning, 2017) and undominated processes, where it could not be determined which processes were driving changes, i.e., neither selection nor dispersal dominated the assembly processes(Jia *et al*., 2020).

**Figure 5:**
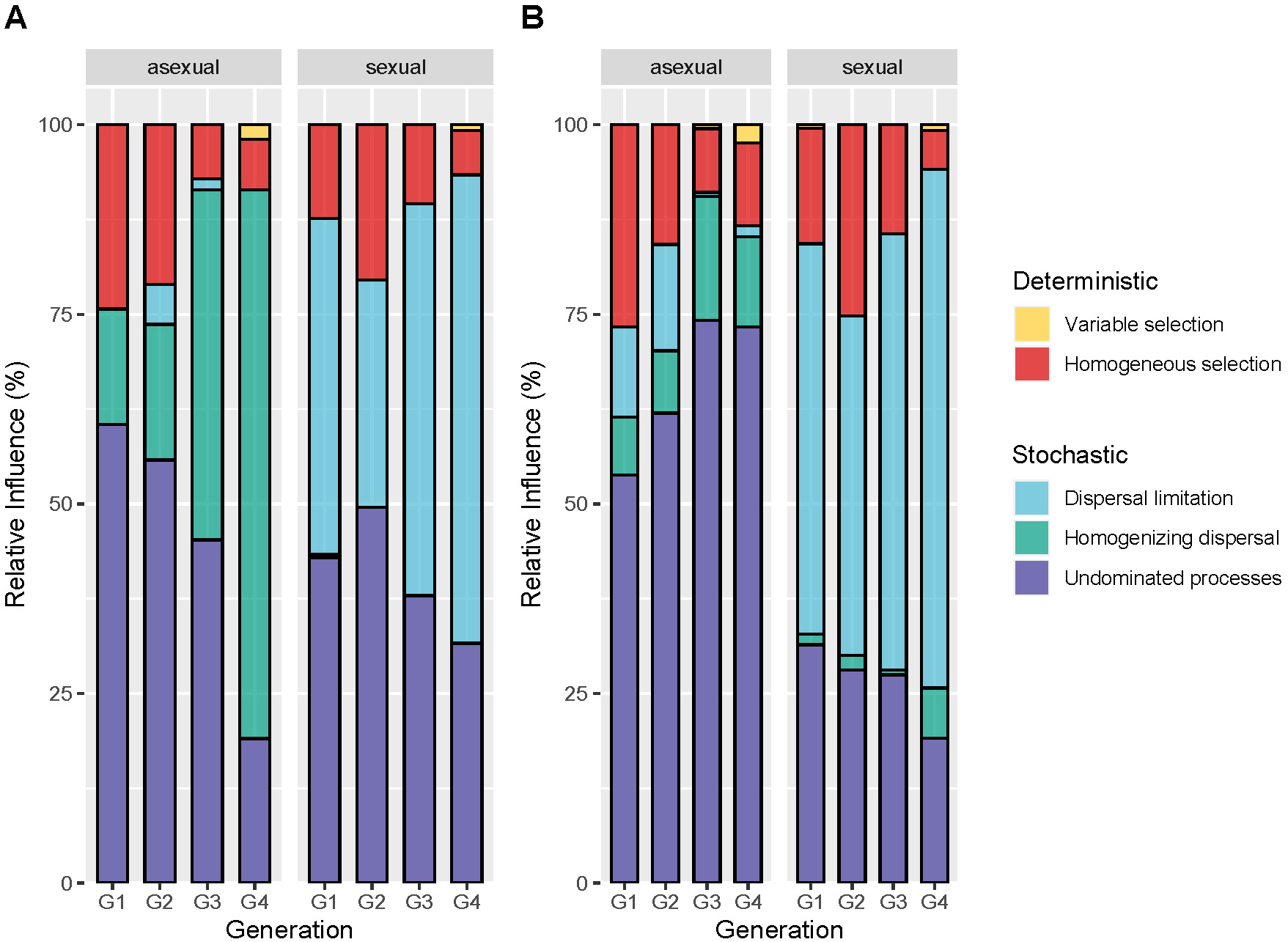
Processes driving bacterial community diversity of seven replicated lines of each asexual (*Wolbachia*-infected) and sexual (*Wolbachia*-uninfected) *A. japonica* line reared over four generations (G1 to G4) in the laboratory in **A)** the full dataset and **B)** after the removal of *Wolbachia* reads. Barplots show the percentage of influencing processes (colours) per generation for the two reproductive modes, asexual (*Wolbachia*-infected) and sexual (uninfected).

Deterministic effects occurred for both reproductive modes (homogeneous selection) and decreased over time (Table S2). In the asexual lines, homogenising dispersal dominated community assembly, next to undominated processes. After generation two, homogenising dispersal was the dominant driver (Fig. 5A; Table S2). The presence of *Wolbachia* reads drove the stochastic process of homogenising dispersal, given that their removal decreased its importance over undominated processes (Fig. 5B). For the sexual lines, dispersal limitation was one of the most critical stochastic processes driving the assembly of bacterial communities, increasing in relative importance over the generations.

## Discussion

Understanding how microbial communities will react to environmental changes is important, given the increasing disturbances of natural habitats by anthropogenic activities. Studying how wild populations change following transfer to controlled laboratory conditions is one way of gaining insight into this. Moreover, investigations of the importance of the microbiome for host fitness, symbiont research and economic rearing often require laboratory experiments in which stable environmental conditions – i.e. a reduced mating pool, and limited scope for horizontal transmission of microbes - induce changes in the host’s microbiome. Here, we investigated the effect of laboratory introduction on bacterial communities associated with an insect host depending on host geographic origin, genetic background (population structure), and the presence of the endosymbiont *Wolbachia,* which modulates host reproduction. As expected, laboratory rearing imposed a reduction in host-associated bacterial diversity. Interestingly, this reduction differed between host reproductive modes (symbiont presence) regarding factors driving the change and timing. This suggests that a powerful symbiont like *Wolbachia*, likely in combination with the host genetic background, plays a significant role in steering alterations in bacterial communities when environmental conditions undergo a shift.

As anticipated, the transition of wasps from their natural habitat into a laboratory setting resulted in decreased bacterial diversity and altered community composition in both sexual and asexual wasps. Indeed, multiple studies found that species reared in the laboratory harbour less diverse microbial communities than their natural counterparts (Gall *et al*., 2017; Waltmann *et al*., 2019; Brown *et al*., 2021). This reduction is likely due to the loss of environmentally obtained microbes from resources such as food and free-living microbes from their environment (Pons *et al*., 2019; Acevedo *et al*., 2021; Luo *et al*., 2021). Interestingly, in our study, this reduction only occurred after the 2^nd^ and 3^rd^ generations for sexual and asexual wasps, respectively. This indicates a maternal transmission of microbes from G0 mothers, who grew up in nature, to their offspring (G1), hatched in the laboratory. Such maternal transmission is well-known in many organisms (Funkhouser and Bordenstein, 2013) but has not been described in *A. japonica*. However, as we did not screen G0 mothers, this awaits further verification.

Our findings also indicate that *Wolbachia* impacted how bacterial communities changed. The delayed reduction in alpha diversity and the gradual homogenisation of the bacterial community composition (beta diversity) in asexual wasps suggests that *Wolbachia* not only shapes the bacterial community of its host but potentially acts as a bacterial community stabiliser (Herren and McMahon, 2018). A similar influence of *Wolbachia* has also been found in fruit flies (*Drosophila melanogaster* (Simhadri *et al*., 2017), the small brown planthopper (*Laodelphax striatellus)* (Duan *et al*., 2020), and artificially infected mosquito adults (*Aedes aegypti)* (Audsley *et al*., 2018). Therefore, the results of our study indicate that the presence of *Wolbachia* may buffer a potential influence of environmental factors affecting the bacterial community of a host (de Vries *et al*., 2004; Russell and Moran, 2006; Ochman *et al*., 2010; Colman *et al*., 2012; Ferguson *et al*., 2018; Duan *et al*., 2020).

We also found that the endosymbiont *Wolbachia* itself is influenced by laboratory introduction. Over four generations, the abundance of *Wolbachia* reads changed to uniformly high levels of *Wolbachia* in wasps from all three asexual lines by the fourth generation. Similar changes in *Wolbachia* were found in *Tetranychus* mites, where some mites experienced an increase in *Wolbachia* titer after laboratory introduction (Zélé *et al*., 2020). Interestingly, our findings indicate that these *Wolbachia* fluctuations are linked to the initial levels of the bacterium. The two northern asexual wasp lines, Kyoto and Sendai (both Pop 5), which initially had a high quantity of *Wolbachia* reads, saw a decline in the endosymbiont after transitioning, followed by a rebound to their initial levels. Conversely, the more southern and genetically distinct Kagoshima population, which initially had a relatively low number of *Wolbachia*, saw an increase in *Wolbachia* over time. The reasons for these initially different *Wolbachia* abundances in the three populations are unknown. However, the subtropical conditions at Kagoshima may have influenced *Wolbachia* abundance as symbionts appear to be sensitive to high temperature (Van Opijnen and Breeuwer, 1999; Hurst *et al*., 2000; Corbin *et al*., 2017; Sumi *et al*., 2017) and show a general geographical pattern of lower prevalence in warmer regions (Corbin *et al*., 2017). Therefore, the more temperate temperatures in the laboratory could have caused an increase.

The found *Wolbachia* fluctuations might have contributed to the decline in bacterial diversity and the shift in microbial community composition by the third generation observed in our data. Potentially, the drop-off in *Wolbachia* numbers at generation two of the two northern lines (Pop 5: Kyoto and Sendai) led to a significant decrease in alpha diversity in their third generation. This may indicate that the stabilising effect of *Wolbachia* was weakened in generation two due to its low abundance. In other words, the impact of *Wolbachia* may not have been strong enough to maintain its control on the bacterial community composition for the next generation. Moreover, the equal abundance of *Wolbachia* in all samples at generation 4 could have been a driver of the homogenisation of the bacterial community composition (beta diversity) of asexual wasps at generation 4. This suggests that symbiont density within an organism is crucial for sustaining symbiont effects, as observed in the correlation between *Wolbachia* density and its protection against viruses in *Drosophila simulans* (Martinez-Sañudo *et al*., 2018). Furthermore, in our study system, *A. japonica*, asexual reproduction relies on a *Wolbachia* threshold (Ma *et al*., 2015), where *Wolbachia* numbers below a certain threshold result in unsuccessful host reproductive manipulation.

Taken together, we find that *Wolbachia* is affected by the introduction into the laboratory with potential knock-on effects on the host-associated microbial community, a decrease in alpha diversity, and a homogenisation of the microbial community composition over time.

Lastly, using multiple replicates per line, we found that stochastic processes mainly drove bacterial community changes for both reproductive modes. This aligns with the theory that host-microbiome variation is predominantly driven by stochastic processes (Adair and Douglas, 2017), as indicated by various studies on community assembly (see (Obadia *et al*., 2017; Vega and Gore, 2017; Sieber *et al*., 2019; Brown *et al*., 2020; Lou *et al*., 2021)). Interestingly, although stochastic processes mainly drove bacterial community changes, we found differences in the processes that dominate changes over generations between the two reproductive modes. In asexual wasps, the process of homogenising dispersal became more dominant over time, likely because the bacterial community became more similar. In contrast, the process of dispersal limitation increased over time in sexual wasps, probably reflecting the higher dissimilarities of the bacterial communities associated with the sexual lines over time. The fact that wasps reacted differently depending on symbiont presence or absence suggests that other factors, apart from environmental factors, must influence bacterial community changes, as all wasps were reared under the same conditions. One of these factors may be the host’s genetic background. Sexually reproducing wasps will likely experience a loss of genetic diversity when introduced to the laboratory due to the reduced genetic mating pool and adaptation to laboratory conditions. This may influence the bacterial community in turn. Indeed, it is known that host genotypes can influence microbial composition, with communities following host phylogenetic signals (Kolasa *et al*., 2019; Lim and Bordenstein, 2020) and having population-specific microbiomes (Bouchon *et al*., 2016; Falony *et al*., 2016; Brinker *et al*., 2019; Rudman *et al*., 2019). Although we did not find clear patterns associated with population structure for alpha and beta diversity of asexual wasps, we cannot exclude that the influence of *Wolbachia* may have overpowered such a signal.

## Conclusion

Here, we showed that microbial communities react differently to environmental changes in the presence of a symbiont. Wasps carrying the endosymbiont *Wolbachia* exhibited a delayed reduction in diversity compared to uninfected wasps. Moreover, this diversity reduction does not always promote homogenisation of the communities, as found for *Wolbachia* infected wasp lines, but can foster their diversification, as seen for uninfected wasps. This suggests that the presence of symbionts with strong host effects, such as *Wolbachia*, can influence how and when a change occurs. Moreover, our results revealed that laboratory introduction reduces host microbial community diversity. This leads to the problem that laboratory-introduced individuals no longer reflect microbiota profiles occurring in nature. Therefore, findings obtained under laboratory conditions regarding microbiota profiles, their function, and the effect on the host, as well as their interaction within the microbial community, may need to be revised or are at least not directly translatable to processes occurring in natural populations. All this highlights the need to bridge the gap from laboratory findings to the effect in the wild. This is a general challenge for researchers and is currently discussed in plant-focused (Sergaki *et al*., 2018; Sessitsch *et al*., 2019) and human-based microbial research (Amato *et al*., 2019) and should be applied to more research fields.

## Data availability statement

Metadata and scripts are available on figshare (10.6084/m9.figshare.29196146). (Files currently under embargo, but can be accessed for reviewers with the following link https://figshare.com/s/54bcd676a21d53add564). Additionally, 16S rRNA data with sample information have been deposited in NCBI’s SRA archives under BioProject ID PRJNA1272023 (accession numbers SAMN48891735–SAMN48891899).

## Acknowledgements, including funding statement and permission to reproduce material from other sources

We thank Fangying Chen for her help in processing the cultures. P.B. thanks Simon Tragust for discussions. P.B. was supported by a scholarship from the Adaptive Life program of the University of Groningen, The Netherlands.

## Ethics statement

All research described in this manuscript was conducted in accordance with the ethical standards of *Environmental Microbiology* and relevant institutional and national guidelines. No human or animal subjects were involved. The authors declare no conflicts of interest and affirm that all data are reported honestly and transparently.

## Author Contributions

**Pina Brinker:** Conceptualisation, Data Curation, Formal Analysis, Investigation, Methodology, Funding Acquisition, Project Administration, Validation, Visualization, Writing – Original Draft Preparation, Writing – Review & Editing; **Joana Falcao Salles:** Funding Acquisition, Writing – Review & Editing, Validation; **Leo W. Beukeboom:** Funding Acquisition, Writing – Review & Editing, Resources; **Michael C. Fontaine:** Funding Acquisition, Writing – Review & Editing, Validation

