## Supplementary information for "*Wolbachia*: The architect of microbial assemblies in response to environmental changes"

Running title: Symbiont-influenced microbiome assembly

Pina Brinker[1,2*§], Joana Falcao Salles[1¥], Leo W. Beukeboom [1$], Michael C. Fontaine[1,3#]

[1] Groningen Institute for Evolutionary Life Sciences (GELIFES), University of Groningen, The Netherlands

[2] Institute of Evolutionary Ecology and Conservation Genomics, University of Ulm, Ulm, Germany

[3] MIVEGEC, Univ. Montpellier, CNRS, IRD, Montpellier, France

[§]ORCID number: https://orcid.org/0000-0003-4723-4077
[¥]ORCID number: https://orcid.org/0000-0003-4317-7263
[$]ORCID number: https://orcid.org/0000-0001-9838-9314
[#]ORCID number: https://orcid.org/0000-0003-1156-4154

**This PDF file includes:**

Table S1-S2

Figures S1-S9

### Tables

**Table S1**: Statistical output of alpha diversity (ASV number and Shannon diversity) analyses for *Wolbachia*-uninfected, sexual (Iriomote, Okinawa, Amami) and *Wolbachia*-infected asexual (Kagoshima, Kyoto, Sendai) lines separately per line (seven replicates per location) once for the full dataset and once for the reduced dataset with *Wolbachia* reads removed. Given are degrees of freedom (DF), F-value (F) and p-value (pVal) of the likelihood-ratio test (LRT) together with the pairwise comparisons between generations (G1 to G4) using Tukey post-hoc test. Sqrt indicates that data needed to be square root transformed.

| Diversity | Location | DF | F | pValue | G1 vs G2 | G1 vs G3 | G1 vs G4 | G2 vs G3 | G2 vs G4 | G3 vs G4 |
|---|---|---|---|---|---|---|---|---|---|---|
| ASV number | Iriomote | 3 | 2.36 | 0.1 | NS | NS | NS | NS | NS | NS |
| ASV number | Okinawa | 3 | 10.26 | <0.001 | <0.001 | <0.001 | 0.0012 | NS | NS | NS |
| ASV number | Amami | 3 | 1.195 | 0.334 | NS | NS | NS | NS | NS | NS |
| ASV number | Kagoshima | 3 | 7.724 | <0.001 | NS | 0.001 | 0.001 | NS | NS | NS |
| ASV number | Kyoto | 3 | 3.277 | 0.041 | NS | NS | NS | NS | NS | NS |
| ASV number | Sendai | 3 | 0.144 | 0.932 | NS | NS | NS | NS | NS | NS |
| Shannon diversiy | Iriomote | 3 | 2.853 | 0.0618 | NS | NS | NS | NS | NS | NS |
| Shannon diversiy | Okinawa | 3 | 4.582 | 0.013 | 0.0268 | NS | 0.0132 | NS | NS | NS |
| Shannon diversiy | Amami | 3 | 2.111 | 0.127 | NS | NS | NS | NS | NS | NS |
| Shannon diversiy | Kagoshima | 3 | 25.86 | <0.001 | NS | <0.001 | <0.001 | <0.001 | <0.001 | NS |
| Shannon diversiy | Kyoto | 3 | 0.563 | 0.646 | NS | NS | NS | NS | NS | NS |
| Shannon diversiy | Sendai | 3 | 1.051 | 0.388 | NS | NS | NS | NS | NS | NS |
| | Without *Wolbachia* | | | | | | | | | |
| ASV number | Iriomote | 3 | 2.01 | 0.143 | NS | NS | NS | NS | NS | NS |
| ASV number | Okinawa | 3 | 8.897 | <0.001 | 0.001 | 0.001 | 0.003 | NS | NS | NS |
| ASV number | Amami (sqrt) | 3 | 1.395 | 0.27 | NS | NS | NS | NS | NS | NS |
| ASV number | Kagoshima | 3 | 2.185 | 0.117 | NS | NS | NS | NS | NS | NS |
| ASV number | Kyoto | 3 | 1.434 | 0.264 | NS | NS | NS | NS | NS | NS |
| ASV number | Sendai | 3 | 0.736 | 0.544 | NS | NS | NS | NS | NS | NS |
| Shannon diversiy | Iriomote | 3 | 2.947 | 0.056 | NS | NS | NS | NS | NS | NS |
| Shannon diversiy | Okinawa | 3 | 4.98 | 0.009 | 0.017 | NS | 0.01 | NS | NS | NS |
| Shannon diversiy | Amami | 3 | 2.4 | 0.094 | NS | NS | NS | NS | NS | NS |
| Shannon diversiy | Kagoshima | 3 | 2.255 | 0.109 | NS | NS | NS | NS | NS | NS |
| Shannon diversiy | Kyoto | 3 | 2.553 | 0.086 | NS | NS | NS | NS | NS | NS |
| Shannon diversiy | Sendai | 3 | 0.487 | 0.696 | NS | NS | NS | NS | NS | NS |

**Table S2**: Percentage table of factors driving changes of asexual (Wolbachia-infected) and sexual (Wolbachia-uninfected) *A. japonica* reared in seven replicated lines over four generations (G1 to G4) in the laboratory in A) the full dataset and B) after the removal of Wolbachia reads. Given are the percentages of deterministic factors (variable selection, homogeneous selection) and stochastic driving forces (dispersal limitation, homogenous dispersal, undominated processes) for each generation per reproductive mode.

**A**

| | With *Wolbachia* | | | | |
|---|---|---|---|---|---|
| | Variable selection % | Homogeneous selection % | Dispersal limitation % | Homogenizing dispersal % | Undominated processes % |
| | | | **Sexuals** | | |
| G1 | 0.00 | 12.38 | 44.29 | 0.48 | 42.86 |
| G2 | 0.00 | 20.48 | 30.00 | 0.00 | 49.52 |
| G3 | 0.00 | 10.46 | 51.63 | 0.00 | 37.91 |
| G4 | 0.74 | 5.88 | 61.76 | 0.00 | 31.62 |
| | | | **Asexual** | | |
| G1 | 0.00 | 24.29 | 0.00 | 15.24 | 60.48 |
| G2 | 0.00 | 21.05 | 5.26 | 17.89 | 55.79 |
| G3 | 0.00 | 7.14 | 1.43 | 46.19 | 45.24 |
| G4 | 1.90 | 6.67 | 0.00 | 72.38 | 19.05 |

**B**

| | No *Wolbachia* | | | | |
|---|---|---|---|---|---|
| | Variable selection % | Homogeneous selection % | Dispersal limitation % | Homogenizing dispersal % | Undominated processes % |
| | | | **Sexuals** | | |
| G1 | 0.48 | 15.24 | 51.43 | 1.43 | 31.43 |
| G2 | 0.00 | 25.24 | 44.76 | 1.90 | 28.10 |
| G3 | 0.00 | 14.38 | 57.52 | 0.65 | 27.45 |
| G4 | 0.74 | 5.15 | 68.38 | 6.62 | 19.12 |
| | | | **Asexual** | | |
| G1 | 0.00 | 26.67 | 11.90 | 7.62 | 53.81 |
| G2 | 0.00 | 15.79 | 14.04 | 8.19 | 61.99 |
| G3 | 0.53 | 8.42 | 0.53 | 16.32 | 74.21 |
| G4 | 2.38 | 10.95 | 1.43 | 11.90 | 73.33 |

#### Figures

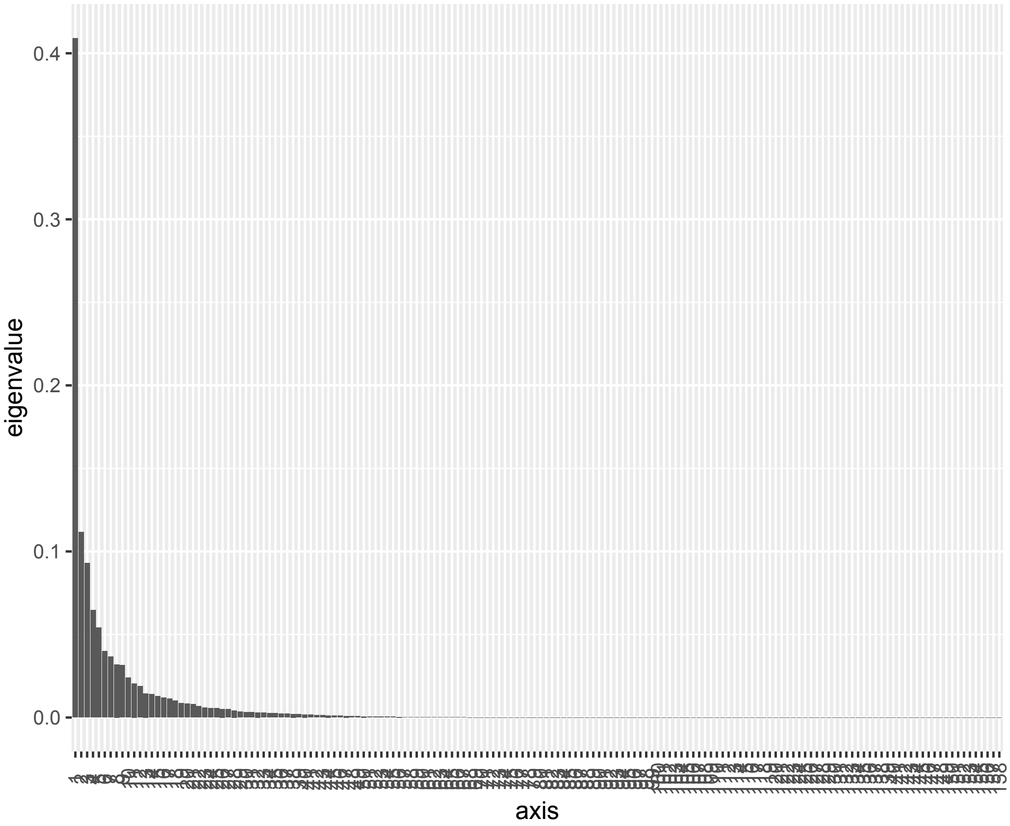


**Fig S1**: Screen plot of principal components vs eigenvalue for the principal coordinates analysis (PCoA) ordination based on the full dataset's Bray–Curtis distance dissimilarity matrices.

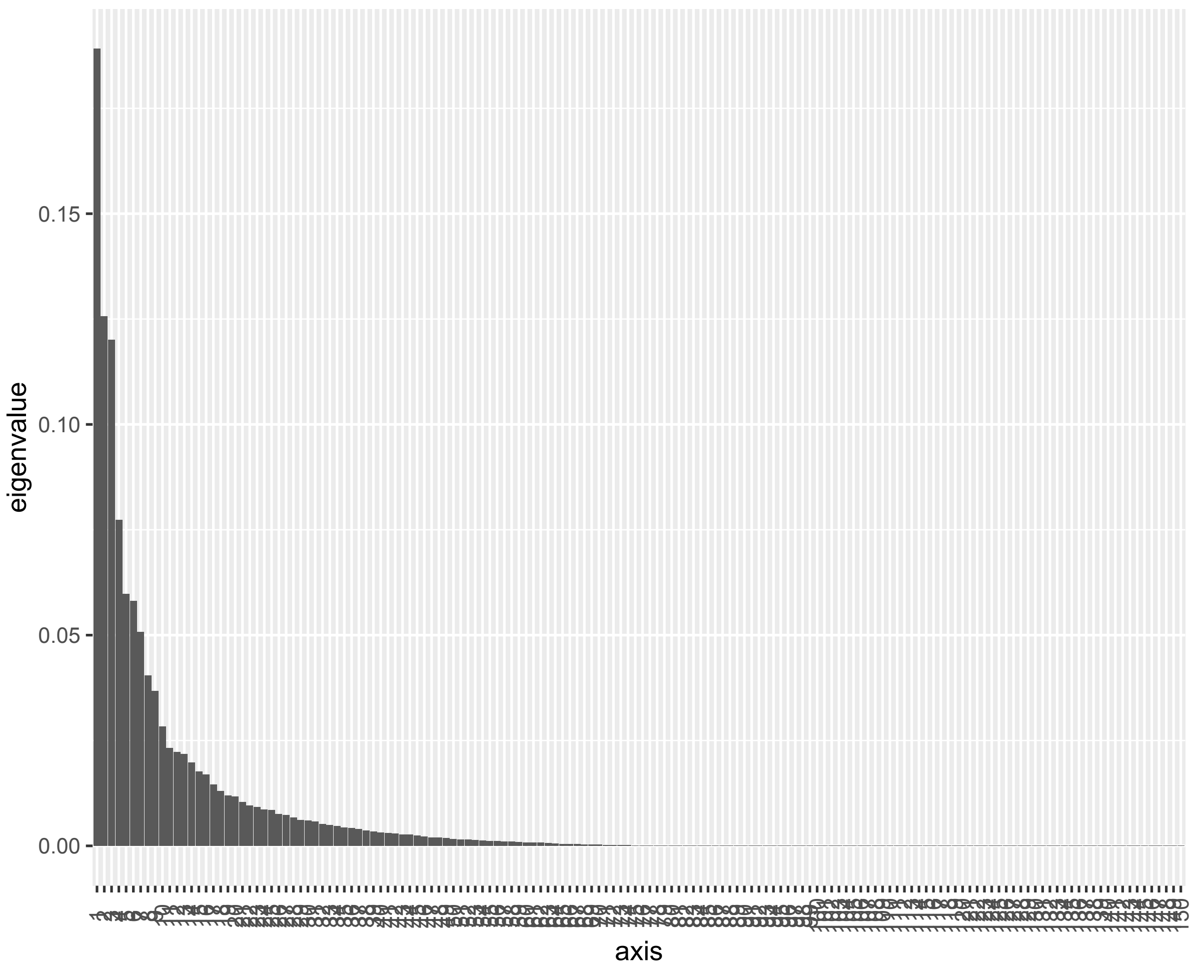


**Fig. S2**: Screen plot of principal components vs eigenvalue for the principal coordinates analysis (PCoA) ordination based on the reduced dataset's Bray–Curtis distance dissimilarity matrices.

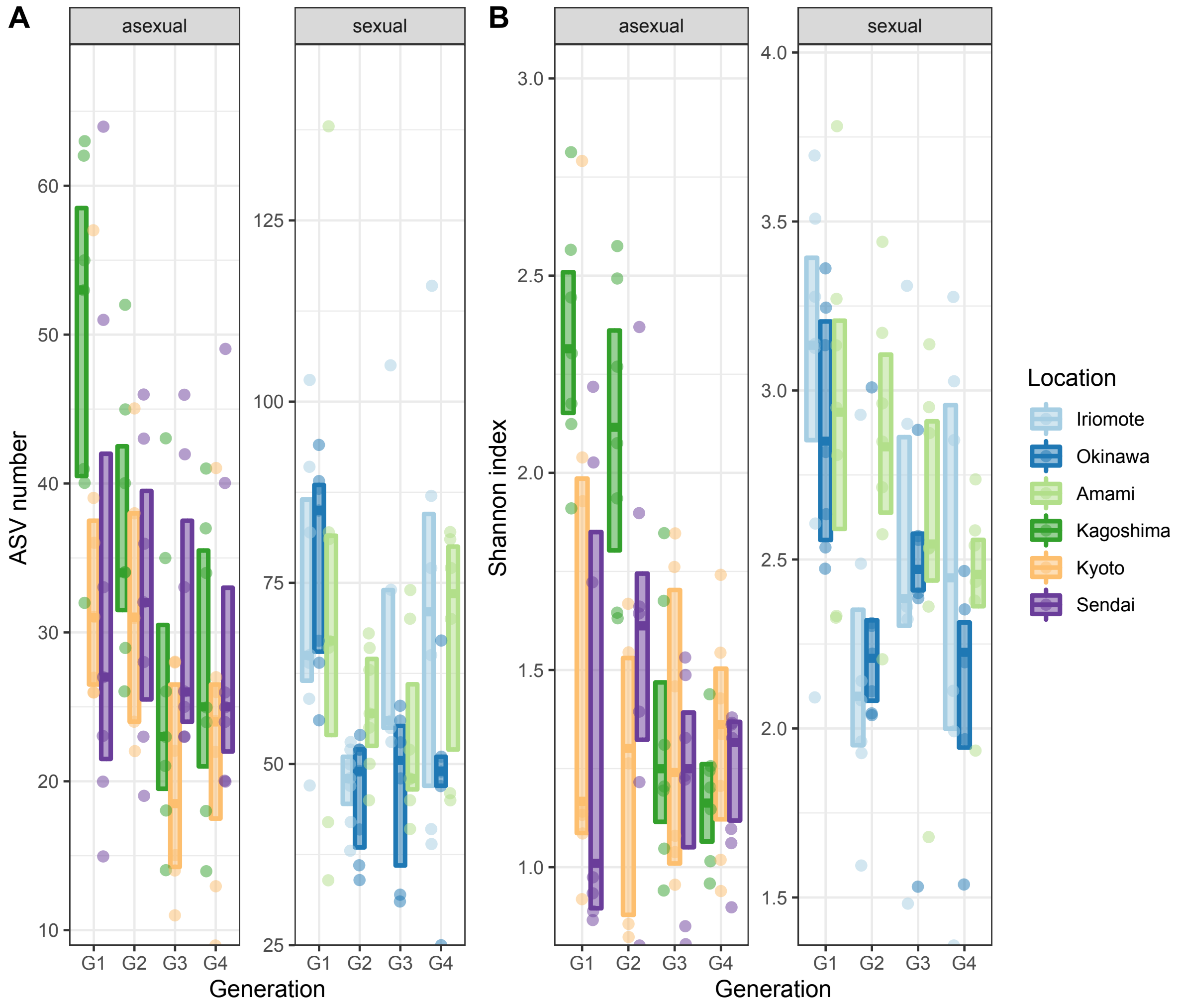


**Figure S3**: Alpha diversity expressed as **A)** ASV number and **B)** Shannon Index of seven replicates of three asexual, *Wolbachia*-infected lines (Kagoshima, Kyoto, Sendai) and three sexual, *Wolbachia*-uninfected lines (Iriomote, Okinawa, Amami) over four generations (G1 to G4) in the laboratory. Only the locations Okinawa and Kagoshima showed statistically significant differences over generations (see Table S1). Boxplots show the median and interquartile range.

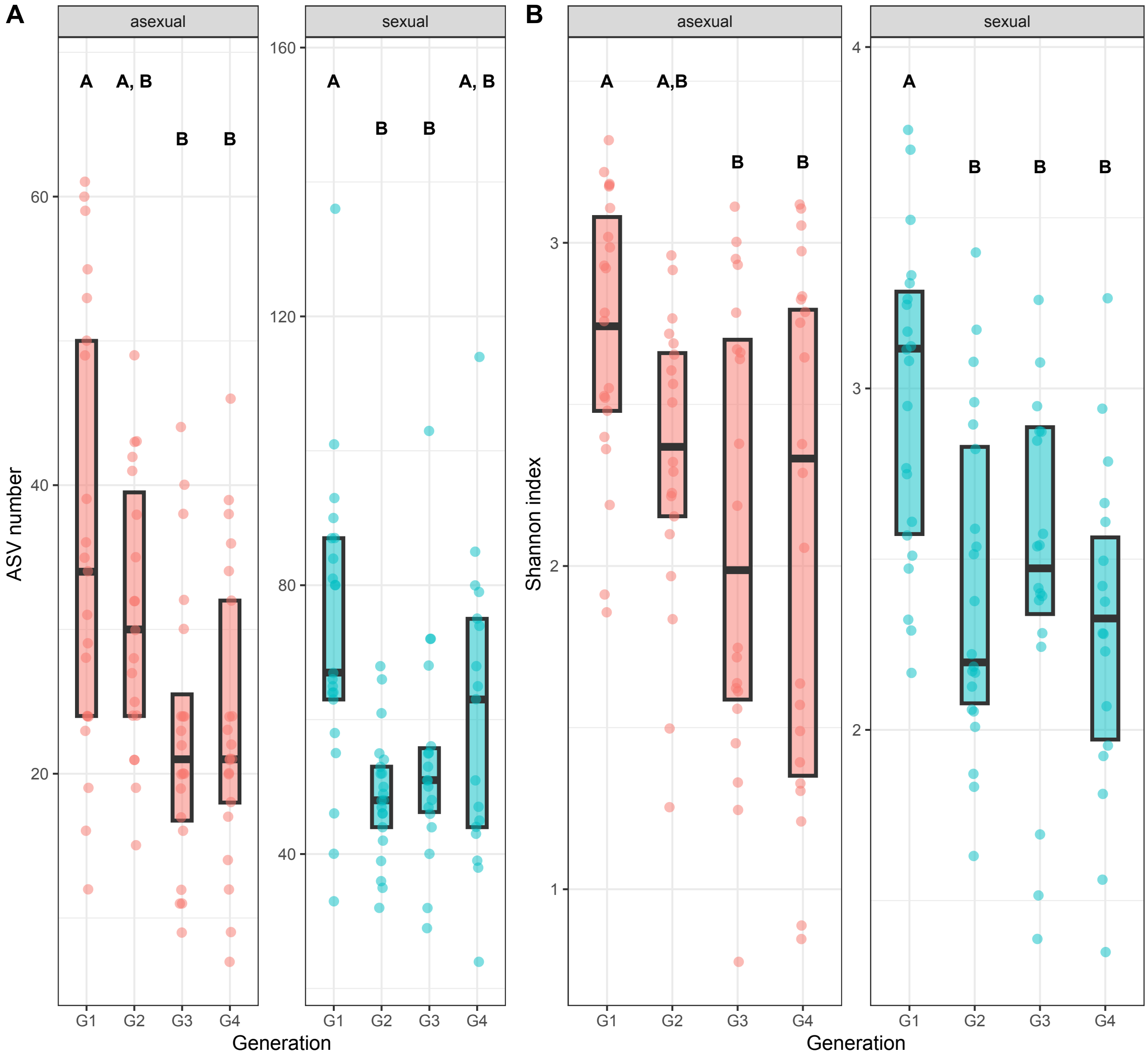


**Figure S4**: Alpha diversity expressed as **A)** ASV number and **B)** Shannon index of asexual (*Wolbachia*-infected) and sexual (*Wolbachia*-uninfected) *A. japonica* wasps reared in seven replicated lines over four generations (G1 to G4) in the laboratory after removal of *Wolbachia* reads from the data. Boxplots show the median and interquartile range. Letters above boxes indicate significant differences between generations.

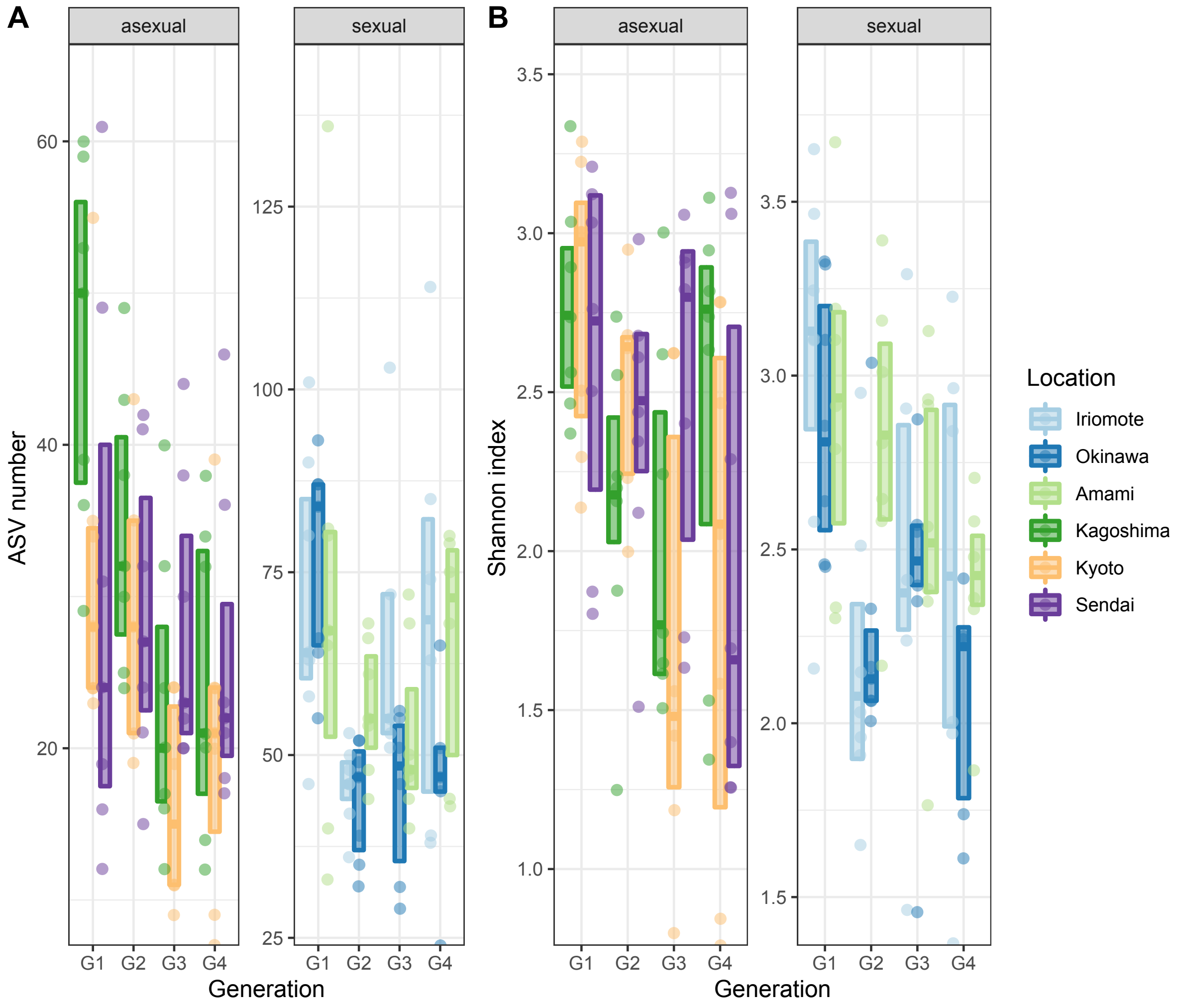


**Figure S5**: Alpha diversity expressed as **A)** ASV number and **B)** Shannon Index of seven replicated lines of three asexual, *Wolbachia*-infected (Kagoshima, Kyoto, Sendai) and three sexual, *Wolbachia*-uninfected lines (Iriomote, Okinawa, Amami) over four generations (G1 to G4) in the laboratory after removal of *Wolbachia* reads from the data. Only the location Okinawa showed statistically significant differences over generations (see Table S1). Boxplots show the median and interquartile range.

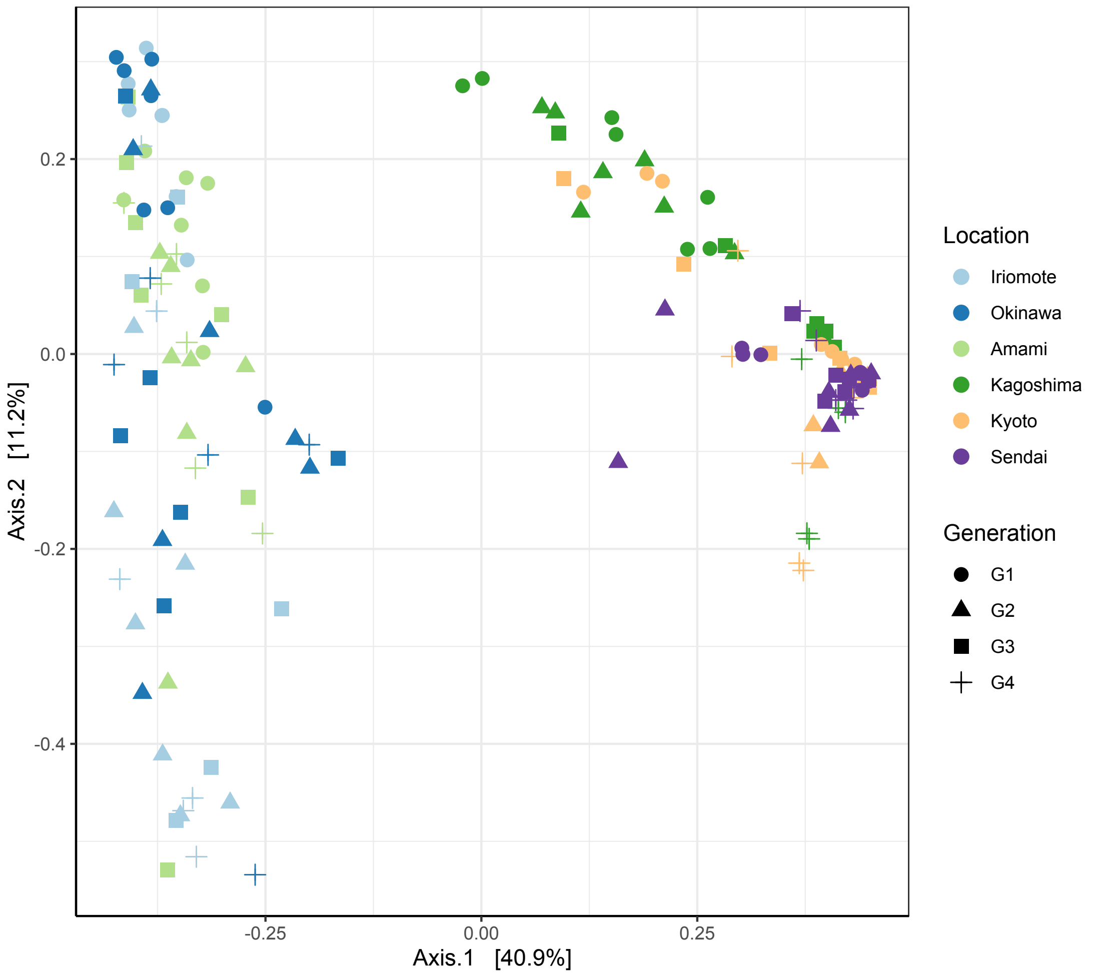


**Figure S6**: Bacterial community composition for the six tested lines of the wasp *A. japonica* over four generations of laboratory culturing. Colours indicate locations, and shapes indicate generation.

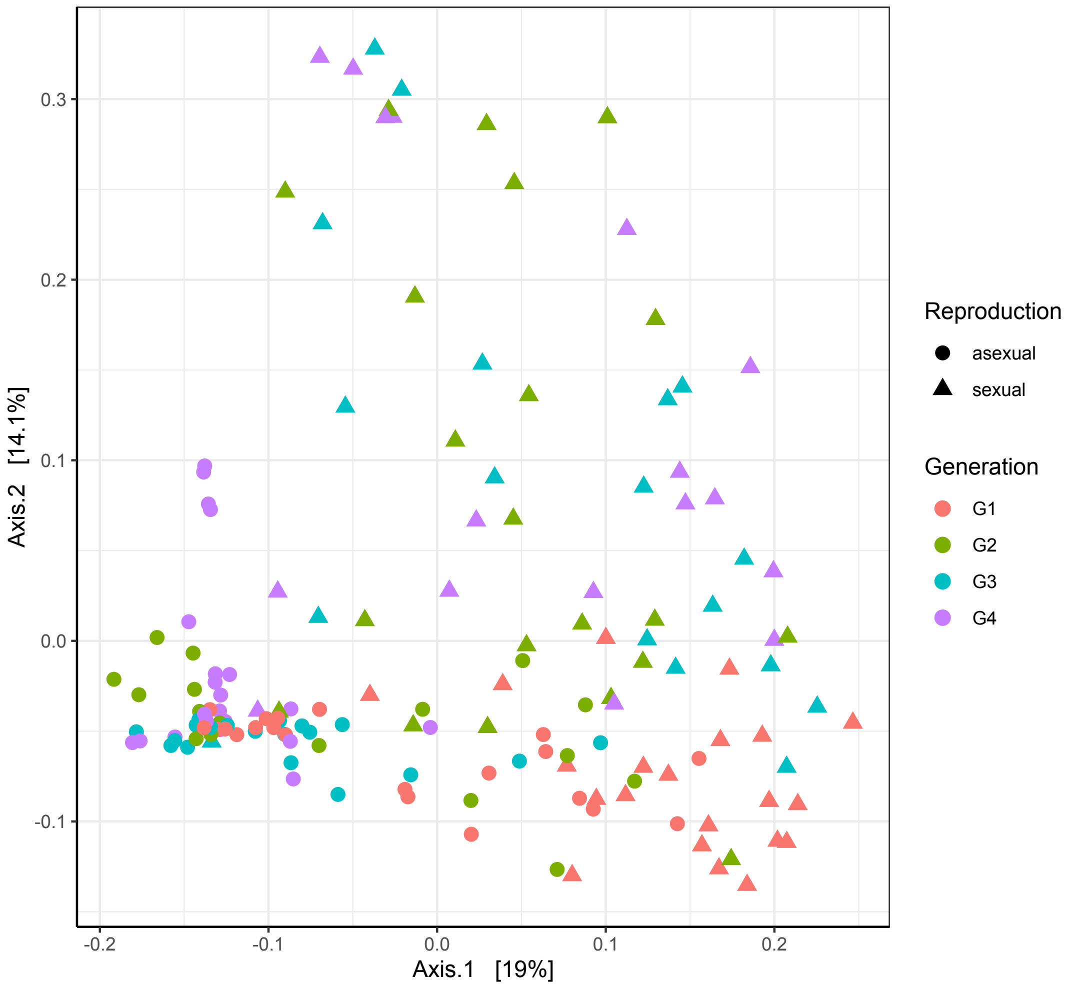


**Figure S7**: Bacterial community composition after removal of *Wolbachia* between the two reproductive modes, sexuals (dots) and asexuals (triangles), of *A. japonica* over four generations of laboratory culturing. Colours indicate locations and shapes generations.

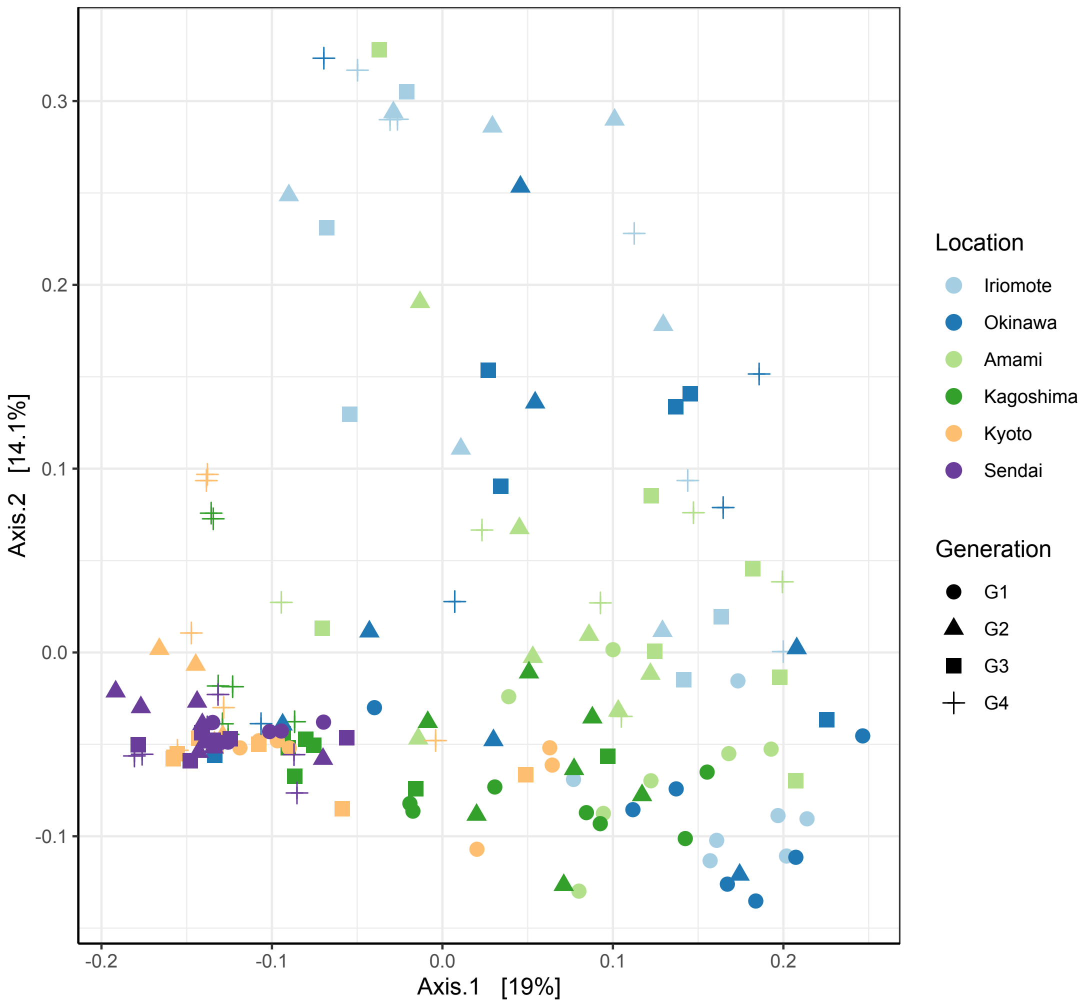


**Figure S8**: Bacterial community composition for the six tested lines of the wasp *A. japonica* over four generations of laboratory culturing after the removal of *Wolbachia*. Colours indicate locations, and shapes indicate generations.

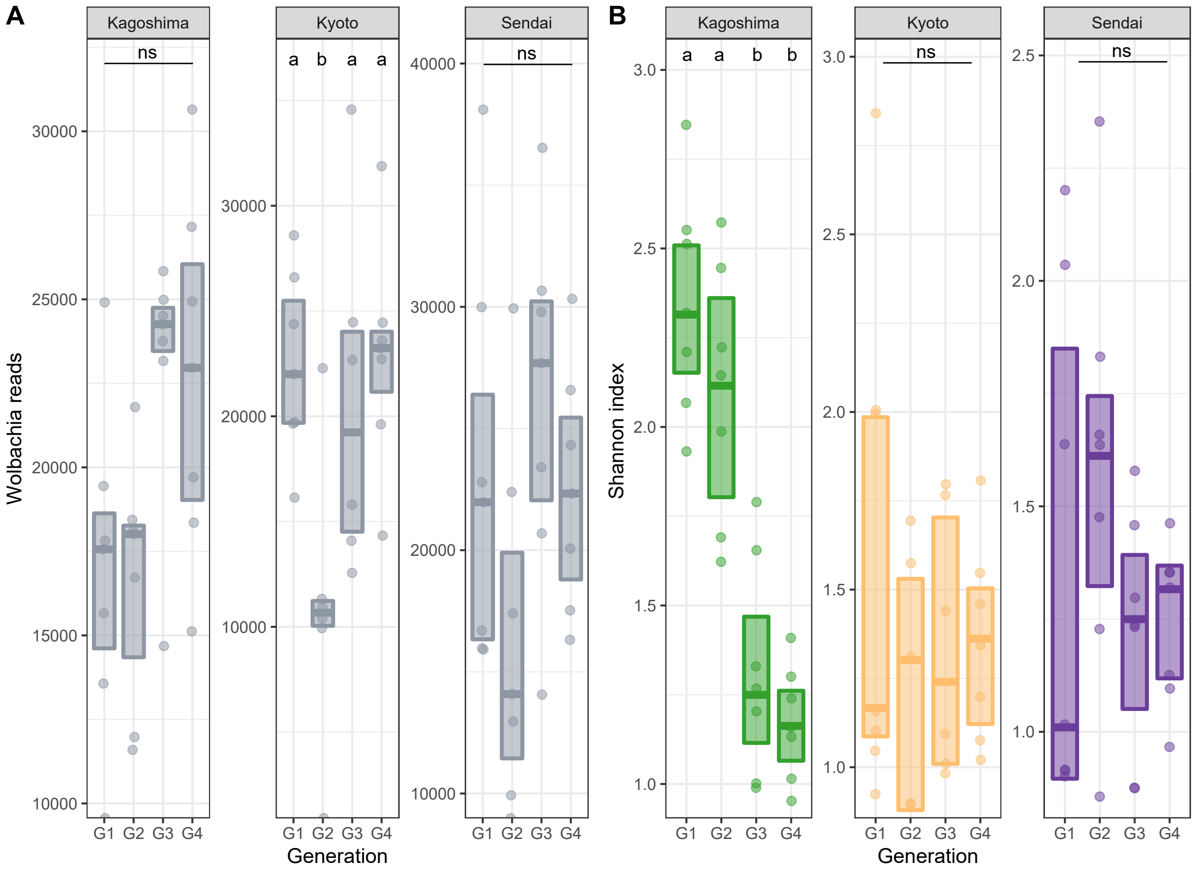


**Figure S9**: **A)** Abundance of *Wolbachia* cells measured as the number of *Wolbachia* reads over four generations (G1-4) from three asexual *A. japonica* lines. Reads extracted out of the unrarefied but normalised dataset are plotted. **B)** Alpha diversity expressed as Shannon Index of seven replicates from three asexual, *Wolbachia*-infected *A. japonica* lines (Kagoshima, Kyoto, Sendai) over four generations (G1 to G4) in the laboratory. Boxplots show the median and interquartile range. Letters above boxes indicate significant differences between generations.
